# In utero exposure to cannabidiol disrupts select early-life behaviors in a sex-specific manner

**DOI:** 10.1101/2022.06.29.498073

**Authors:** Daniela Iezzi, Alba Caceres, Pascale Chavis, Olivier J.J. Manzoni

## Abstract

Cannabidiol (CBD), one of the main components of cannabis, is generally considered safe, despite the lack of studies on the possible consequences of its consumption during critical periods of neurodevelopment, including prenatal life. Although CBD crosses the placenta and its use during pregnancy is steadily increasing, the impact of gestational CBD exposure on early life is unknown. Here, we combined behavioral exploration and deep learning to assess how *in utero* exposure to low doses of CBD alters pre-weaning behaviors in mouse pups of both sexes. The data reveal that pups from CBD-treated dams exhibit sex-specific alterations in weight growth, homing behavior, and the syllabic repertoire of ultrasonic vocalizations. Thus, prenatal CBD is associated with alterations in innate behavioral responses and communication skills.

## Introduction

While Δ9-THC is the component of most concern in *Cannabis sativa* L. in terms of prenatal exposure, the plant contains over 300 compounds, including cannabidiol (CBD). Although structurally similar, CBD does not induce the psychotropic effects classically associated with Δ9-THC (1,2). Consequentely, CBD is globally perceived as safe and free of harmful side effects. Its clinical interest is due to its potential benefits as a natural antipsychotic, anti-nociceptive, anticonvulsant, antiemetic, anxiolytic, anti-inflammatory, antioxidant, and neuroprotective agent (3– 6). Despite the lack of scientific evidence regarding safety of CBD during gestation, pregnant women use CBD for a plethora of pregnancy-related symptoms including nausea, insomnia, anxiety, and chronic pain (7). CBD crosses the placenta and alters its very structure, both of which can have a significant impact on pregnancy outcomes (8–10). Moreover, a recent study showed that extended exposure of CD1 mice to CBD spanning from gestation through the first week after birth alters repetitive and hedonic behaviors in the adult progeny (11). The paucity of preclinical data on the impact of *in utero* CBD exposure prompted us to investigate the postnatal impact of gestational CBD exposure to assess potential risks associated with CBD use during this period. The data reveal that pups from CBD-treated dams exhibit previously unknown sex–specific cognitive alterations in early-life.

## Materials and Methods

### Animals

Male and female C57BL6/J (8-10 weeks age) were purchased from Janvier Lab and housed in standard wire-topped Plexiglas cages (42 cmx27 cmx14 cm), in a temperature and humidity-controlled condition (i.e., temperature 21 ± 1°C, 60 ± 10% relative humidity and 12 h light/dark cycles). Food and water were available ad *libitum*. After one week of acclimation, the female pairs were placed with a single male mouse in the late afternoon. The morning the vaginal plug was found was designated as day 0 of gestation (GD0) and pregnant mice were housed individually. From GD5 to GD18, dams were injected subcutaneously (s.c.) daily with vehicle or 3 mg/Kg of CBD (Nida Drug Supply Program), dissolved in a vehicle consisting of Cremophor EL (Sigma-Aldrich), ethanol, and saline at 1:1:18 ratios, and administered at volume of 4 mL/Kg. Control dams (SHAM) were injected the same volume of vehicle solution. This dose of CBD reaches the embryonic brain and cause some behavioral changes in the offspring (11). For each litter, the date of birth was designated as postnatal day (PND) 0. The behavioral tests were performed in male and female offspring during perinatal period (PNDs 10 and 13). Body weight of SHAM and CBD pups was measured every 3 days until one day after weaning (PND 22). All procedures were performed in conformity with the European Communities Council Directive (86/609/EEC) and the United States NIH Guide for the Care and Use of Laboratory Animals. The French Ethical committee authorized the project APAFIS#3279.

### Behavioral tests

#### Ultrasound vocalizations (USVs)

USVs were induced by quick maternal separation in male and female pups at PND 10 as previously described (12). Each tested mouse was placed into an empty plastic container (11 × 7 × 3.5 cm), located inside a sound-attenuating isolation box (32 × 21 × 14 cm). USVs were recorded using an ultrasonic microphone (Ultravox Noldus), connected via the Ultravox device (Noldus, Netherlands) and placed 20 cm above the pup in its plastic container. At the end of 4-minute recording session, each pup was weighed, and the body temperature checked.

Acoustic analysis was performed using DeepSqueak (Version 2.6.2), a deep learning-based software for the detection and analysis of USV (for more details see, (13)). The audio file was individually transferred into DeepSqueak, converted in the corresponding sonograms, and analyzed using a Faster-RCNN object detector. The lower and higher cut-off frequency, 20 kHz and 100 kHz respectively, were applied in order to reduce the background noise outside the relevant frequency. Once detected, each sonogram was converted in the corresponding spectrogram and the calls identified as noise were manually removed. Call classification, transitional probabilities and syntax analysis were performed automatically with DeepSqueak’s built in mouse call classification neural network. USV parameters were classified based on a quantitative and qualitative analysis. The *quantitative* analysis included the percentage of vocalizing and non-vocalizing mice in each group, the number of calls, the latency to the first vocalization (in sec.), the mean call duration (in msec), and the mean dominant frequency (in kHz). Whereas the *qualitative* analysis focused on the study of vocal repertoire based on syntax analysis (i.e., different categories of calls) and the transitional probabilities for each group. The latter was expressed as the probability that one type of call followed the previous one and the following calls were indicated on the x-axis.

#### Homing test

The homing test was performed as published (14), in SHAM and CBD mice previously tested for the USVs. At PND 13 both male and female pups were separated from the dam, and kept for 30 min in a different cage on a heating pad set at the temperature of 35°C. Each tested mouse was placed in the Plexiglas cage (21x 15 cm) which had one-third of the litter from the pup’s original cage and two-thirds of clean litter. The latter was considered as the unfamiliar area, while the one with the old litter was the nest area. The pup was located at the edge of the clean bedding and its behavior was videorecorded for the following 4 min. Homing performance was performed using Ethovision and considering the locomotory activity (in cm.), the velocity (in cm/sec), the moving time (in cm), the latency to reach the nest (in sec.) and the distance moved (in cm), the time spent (in sec.) and the entries in the nest and in the unfamiliar area.

### Statistical analysis

Statistics were performed with GraphPad Prism 9 and DeepSqueak 2.6.2. Behavioral data were tested for the Normality and statistical analysis was performed with Multiple Mann-Whitney *U* test. N values corresponds to the number of animals tested for each group. When achieved, the significance was expressed as exactly *p*-value in the figures. The ROUT test was applied to all data sets to identify outliers, which were then excluded from the data sets.

## Results

### Gestational CBD affect postnatal growth in a sex specific manner

Prenatal cannabis exposure influences neonatal outcome in multiple ways (15) and preclinical data indicate that gestational THC reduces body weight in early life. In contrast, the effects of *in utero* CBD exposure are unknown. Dams were given a low dose of CBD (3 mg/Kg, s.c.) or a vehicle (SHAM) once daily from GD5 to GD18 and their pups’ body weights were monitored throughout the perinatal period until weaning (PND 10-22; for more details see Table 1). Body weights of pups from CBD-exposed dams were consistently higher than those of SHAM dams (Fig. 1). Remarkably, the increase in body weight was observed exclusively in male offspring (Fig. 1A), whereas the weight of *in utero* exposed females was indistinguishable from that of SHAM females (Fig. 1B). Thus, gestational CBD alters the growth trajectory in a sex-specific manner.

**Table 1:** Cannabidiol prenatal exposure increases pup weights in a sex-specific way. Male and female pup weights were collected every three days from postnatal day 10 to postnatal day 22. Values are expressed as median, maximum and minimum. The *p* values are given for each day as compared with pups from sham-treated dams on the same postnatal day, as determined by Multiple Mann-Whitney *U* test.

| Postnatal day weights (g) | TREATMENT | Median | Max | Min | N | *p* value<br>Multiple Mann-Whitney<br>Unpaired t test |
| --- | --- | --- | --- | --- | --- | --- |
| PND 10 | SHAM MALE | 3.4 | 6.7 | 3.1 | 18 | 0.003 |
|  | CBD MALE | 5.5 | 5.8 | 4 | 14 |  |
|  | SHAM FEMALE | 3.5 | 5.9 | 3.1 | 14 | 0.06 |
|  | CBD FEMALE | 5.1 | 5.8 | 3.9 | 12 |  |
| PND 13 | SHAM MALE | 4.9 | 8 | 3.6 | 18 | 0.003 |
|  | CBD MALE | 6.6 | 7.4 | 5.7 | 14 |  |
|  | SHAM FEMALE | 5 | 8.6 | 3.5 | 14 | 0.08 |
|  | CBD FEMALE | 6.4 | 7.3 | 5.6 | 12 |  |
| PND 16 | SHAM MALE | 6 | 9.4 | 4.2 | 18 | 0.04 |
|  | CBD MALE | 7.2 | 8.2 | 6.3 | 14 |  |
|  | SHAM FEMALE | 6 | 9.6 | 3.9 | 14 | 0.27 |
|  | CBD FEMALE | 7 | 8.2 | 6.1 | 12 |  |
| PND 19 | SHAM MALE | 6.9 | 9.7 | 5 | 18 | 0.01 |
|  | CBD MALE | 8.45 | 9.9 | 6.6 | 14 |  |
|  | SHAM FEMALE | 7.3 | 10 | 4.2 | 14 | 0.09 |
|  | CBD FEMALE | 8.2 | 9.8 | 6.2 | 12 |  |
| PND 22 | SHAM MALE | 8.2 | 11.8 | 5.7 | 18 | 0.002 |
|  | CBD MALE | 10.5 | 11.8 | 7.5 | 14 |  |
|  | SHAM FEMALE | 7.9 | 11 | 5.6 | 14 | 0.20 |
|  | CBD FEMALE | 8.6 | 10.6 | 6.8 | 12 |  |

**Figure 1.**
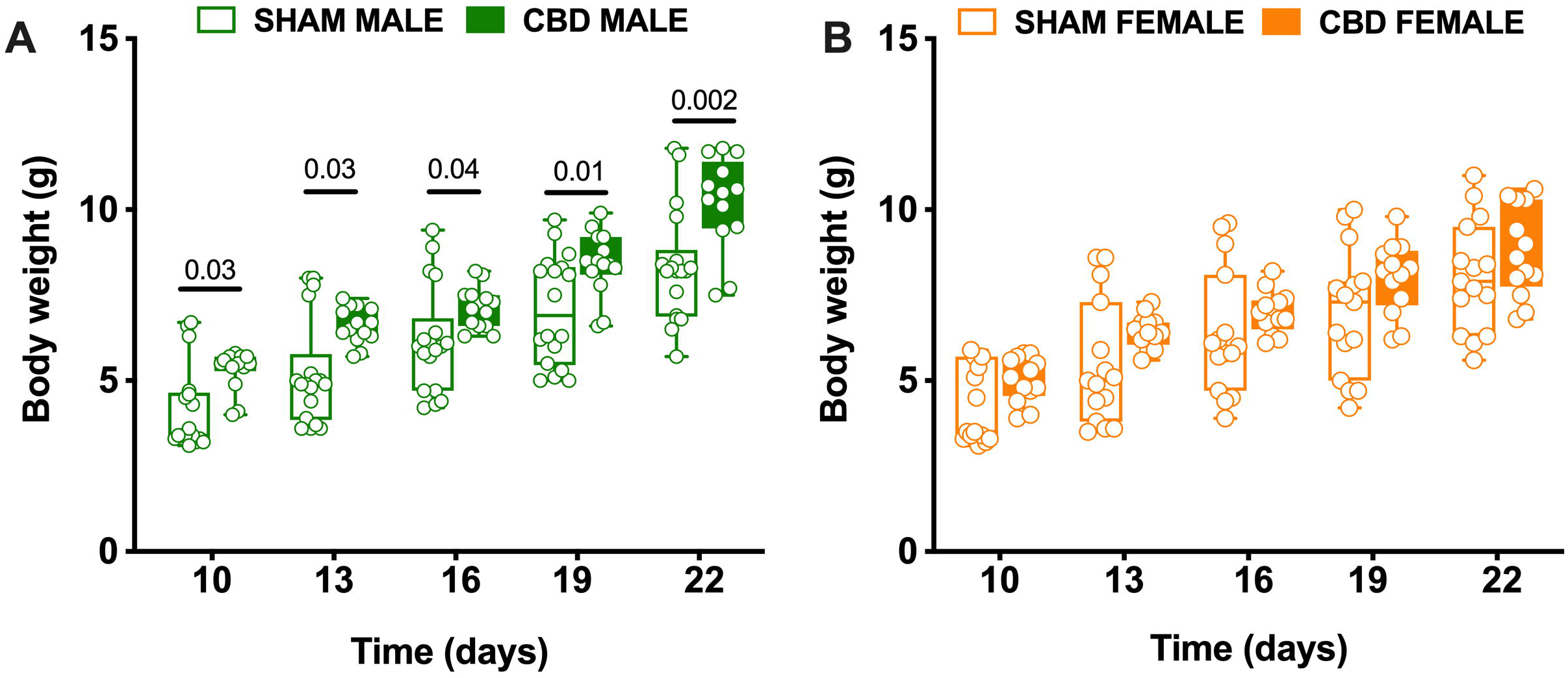
Prenatal exposure to CBD specifically increases the body weight of male offspring. (**A-B**) Starting at PND 10, SHAM and CBD pups of both sexes were weighed every 3 days until one day after weaning (i.e., 22). Fetal CBD exposure was associated to increase of bodyweight in male (**A**) but not female progeny (**B**), (SHAM MALE N = 18 light green, CBD MALE N = 14 dark green, SHAM FEMALE N = 14 light orange, CBD FEMALE N = 12 dark orange). Data are represented as Box and whisker plots (first quartile, median, third quartile). Individual data point represents a single animal. Multiple Mann-Whitney *U* test.

### Gestational CBD modifies the coarse characteristics of ultrasonic communications in a sex-specific manner

Offspring-mother communication is necessary for mouse pups, who emit ultrasonic vocalizations (USV) to convey their emotional conditions (16). Thus, upon separation from their mother and nest, rodent pups vocalize to engage maternal care (17). Perinatal cannabinoids (i.e., Δ9-THC or cannabimimetics) negatively impact neurodevelopment (18–21) and strongly alter USV emissions (22,23). We first quantified the coarse characteristics (i.e., number, latency, duration, and frequency) of separation-induced calls emitted by pups in our different groups (Fig. 2 and Table 2). Prenatal CBD altered the proportion of vocalizing and non-vocalizing pups: more males in the CBD-exposed groups did not vocalize at all during the recording session compared with all other groups (Fig. 2A-B). The total number and latency of first vocalization were similar across treatments and sex (Fig. 2C-D). However, marked differences in the sex-specific effects of prenatal CBD were evident in the mean duration and mean frequency of USVs (Fig. 2E-F). CBD males made shorter calls (Fig. 2E), whereas CBD females made more high frequency calls (Fig. 2F) than their SHAM counterparts. The probability distribution of USV frequency followed a bi-modal distribution in CBD-exposed males but not in females (Fig. 2G-H). Finally, a minor mode corresponding to high frequency calls was specifically observed in CBD-exposed males (Fig. 2G).

**Table 2:** Fetal CBD modifies the frequency and the duration of perinatal USVs. Data were collected from litters for each condition as described in Methods and Materials. Values are expressed as median, maximum and minimum. Significant difference was observed in the Mean USVs duration in CBD compared to SHAM male pups and in the Mean frequency duration in CBD compared to SHAM female pups as determined by Multiple Mann-Whitney *U* test.

| Parameter | PND | TREATMENT | Median | Max | Min | N | *p* value<br>Multiple Mann-Whitney<br>Unpaired t test |
| --- | --- | --- | --- | --- | --- | --- | --- |
| Number of USVs | 10 | SHAM MALE | 67 | 258 | 3 | 14 | 0.1052 |
|  |  | CBD MALE | 31 | 96 | 1 | 10 |  |
|  |  | SHAM FEMALE | 77 | 320 | 4 | 11 | 0.0785 |
|  |  | CBD FEMALE | 18 | 130 | 2 | 11 |  |
| Latency (sec) | 10 | SHAM MALE | 4.42 | 163.38 | 0.01 | 14 | 0.0821 |
|  |  | CBD MALE | 32.56 | 256.39 | 1.34 | 10 |  |
|  |  | SHAM FEMALE | 20.29 | 56.35 | 0.32 | 11 | 0.26 |
|  |  | CBD FEMALE | 38.64 | 212.51 | 0.07 | 11 |  |
| Mean USVs duration (sec) | 10 | SHAM MALE | 0.05 | 0.09 | 0.02 | 14 | 0.03 |
|  |  | CBD MALE | 0.03 | 0.06 | 0.02 | 10 |  |
|  |  | SHAM FEMALE | 0.05 | 0.08 | 0.02 | 11 | 0.6396 |
|  |  | CBD FEMALE | 0.04 | 0.08 | 0.01 | 11 |  |
| Mean Frequency (KHz) | 10 | SHAM MALE | 56.05 | 65.00 | 51.81 | 14 | >0.9999 |
|  |  | CBD MALE | 56.15 | 67.11 | 47.08 | 10 |  |
|  |  | SHAM FEMALE | 56.47 | 63.46 | 53.39 | 11 | 0.004 |
|  |  | CBD FEMALE | 59.76 | 64.12 | 56.21 | 11 |  |

**Figure 2.**
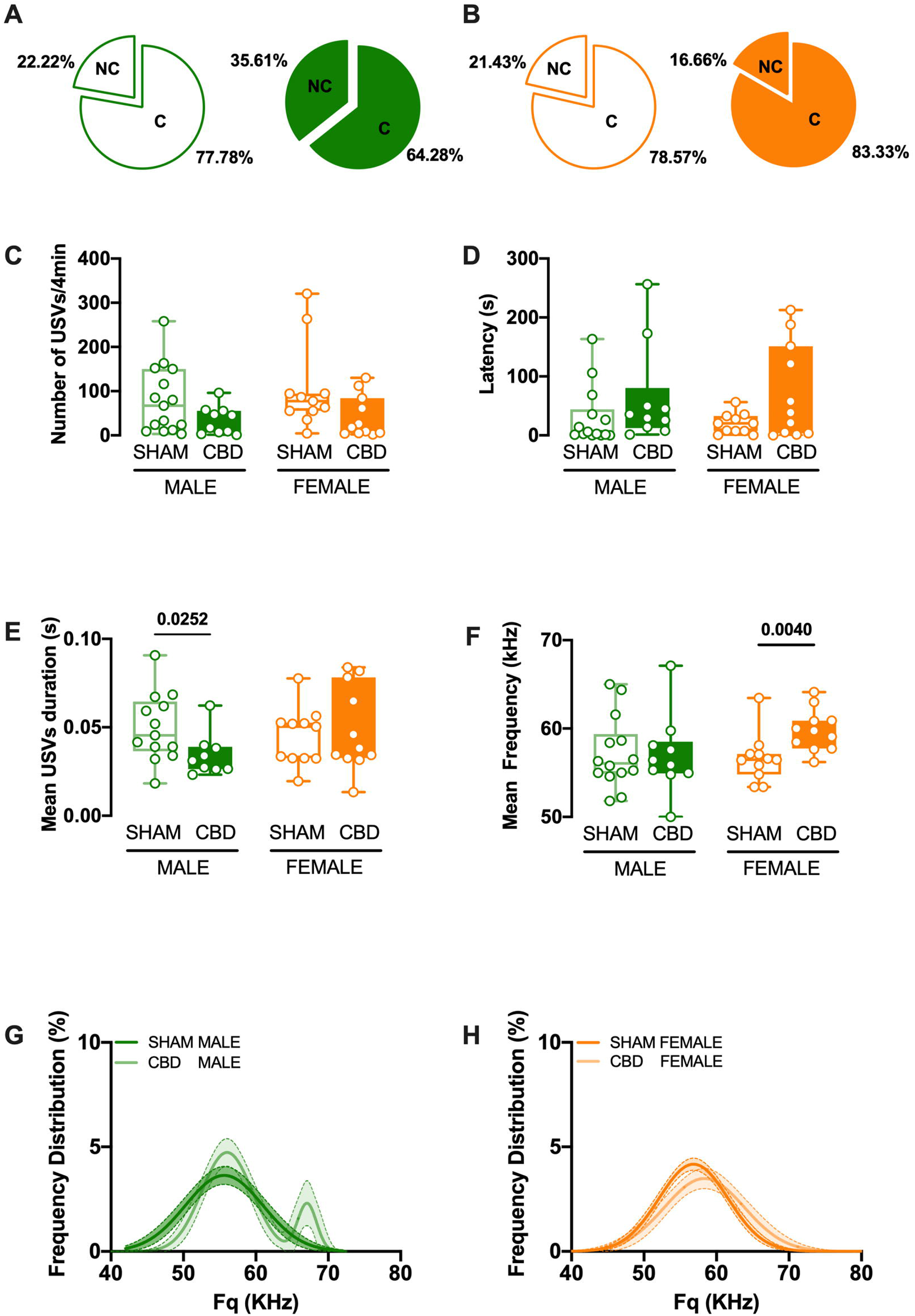
Fetal CBD modifies the frequency and the duration of perinatal USVs. (**A-B**) Pie graphs depicting the percentages of vocalizing (C: call) and non-vocalizing (NC: no call) mice, in SHAM and CBD male (left) or female (right) pups. Percentages were calculated as number of animals vocalizing or not / total number of tested animals. Quantitative analysis of USVs shows that following CBD prenatal exposure, the total number of USVs (**C**) and the latency to the first vocalization (**D**) were similar to those of sham, in both sexes. In contrast, mean duration (**E**) and mean frequency (**F**) of emitted calls were modified in a sex-specific manner in the CBD progeny. The mean call duration was shorter in CBD male, while the mean frequency was higher in CBD females, compared to their respective SHAM group, (SHAM MALE N = 14 light green, CBD MALE N = 10 dark green, SHAM FEMALE N = 11 light orange, CBD FEMALE N = 11 dark orange). Data are represented as Box and whisker plots (first quartile, median, third quartile). Individual data points represent a single animal. Multiple Mann-Whitney *U* test, *p<0.05. (**G**) The average distribution of call frequency was monomodal (i.e., fitted with a single Gaussian function, dark green) in SHAM male, but bimodal (i.e., fitted by the sum of two Gaussians for CBD male, light green). (**H**) In contrast, the call frequency was monomodal in SHAM (light orange) and CBD exposed female pups (dark orange). Data are represented as curve fit (±CI) of the principal frequency’s average distribution.

### Prenatal CBD modifies the syllabic repertoire of ultrasonic communication in a sex-specific manner

Does prenatal CBD alter vocalization patterns in pups? To address this question, we analyzed the amount and spectral characteristics of USVs detected in our different groups with DeepSqueak (13). We first compared the vocal repertoire of SHAM and CBD pups of both sexes. Call categorization (Fig. 3 and Table 3) showed that, while the number of USVs was similar, CBD gestation largely affected the vocal repertoire (Fig. 3). Thus, the proportion of each type of call made during the 5-min test was compared in male and female CBD pups and their control counterparts (Fig. 3B-C-E-F). Notably, male and female CBD pups showed a significant reduction in Complex Trill, Downward Ramp, and Inverted-U calls (Fig. 3D, G) compared with SHAM pups. Furthermore, only CBD male pups showed a significant increase in Short, Trill, and Step-up call (Fig. 3D). In addition, we observed that CBD females emitted significantly less Flat calls compared to SHAM females (Fig. 3G). Thus, call syntax analysis showed multiple sex-specific differences in the vocal repertoire of CBD-exposed pups.

**Figure 3.**
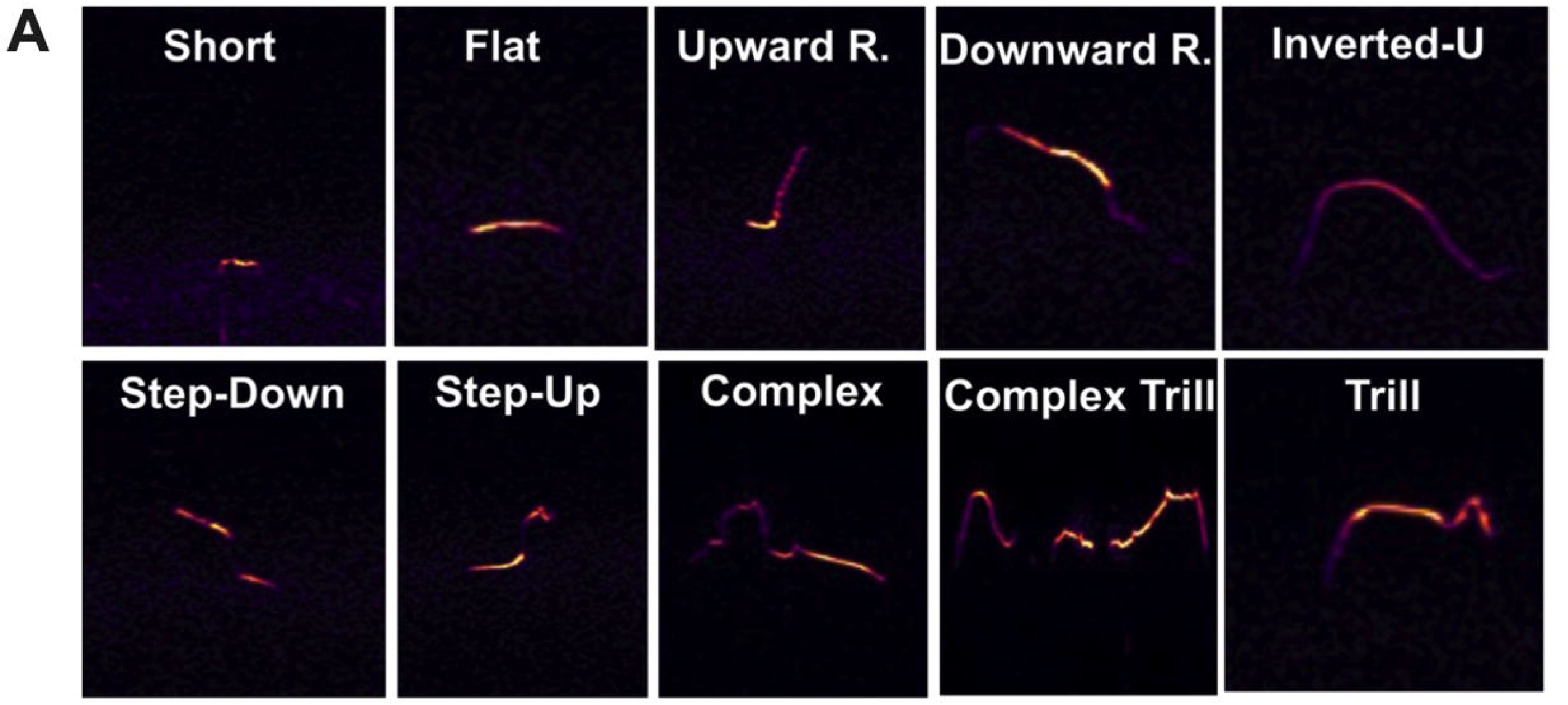
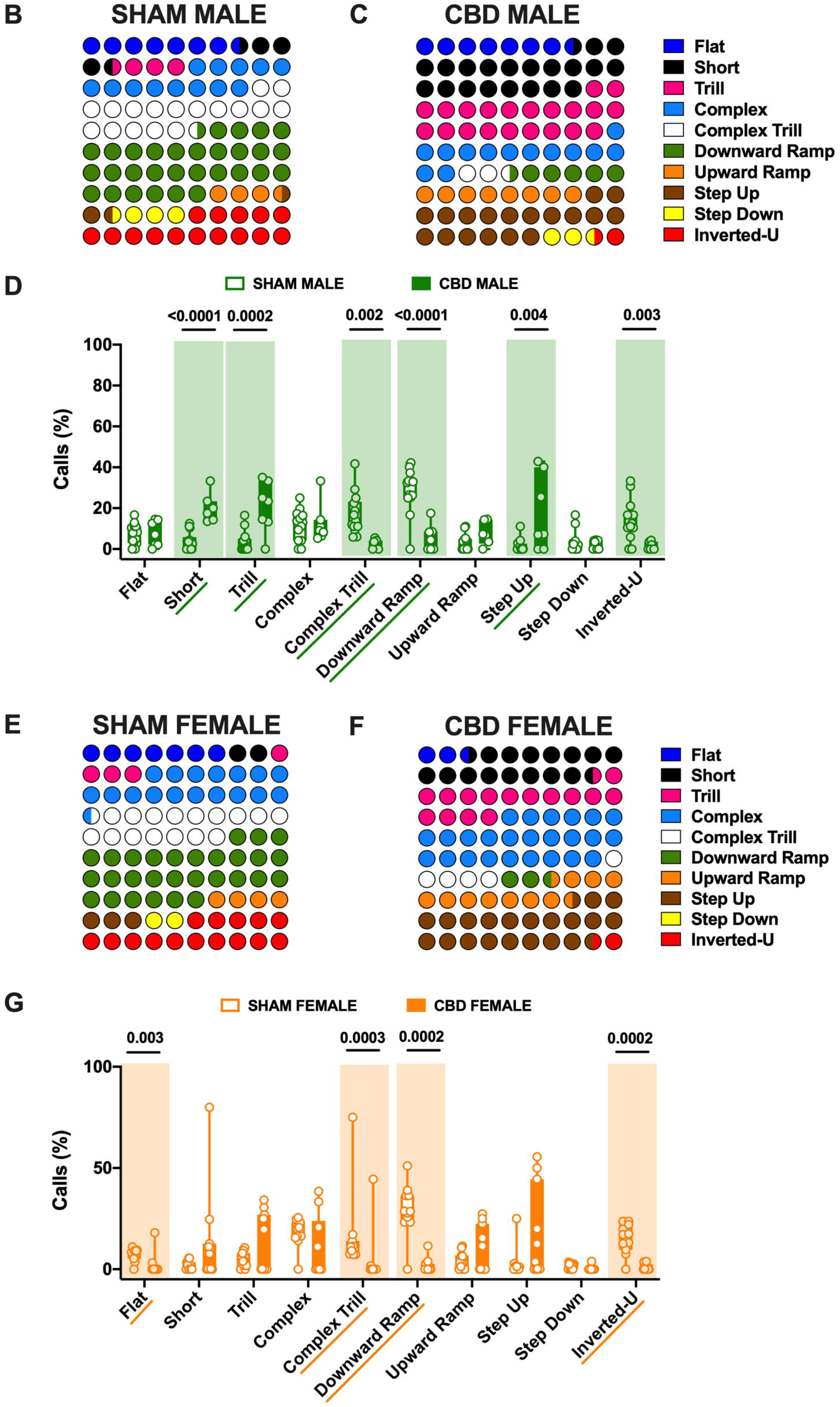
Call type profile is altered in a sex-specific manner in CBD-exposed pups. (**A**) Representative USV calls classified into ten distinct categories based on Supervised-call classification neural network (DeepSqueak). Color maps of USVs showing differential distributions of call types emitted by SHAM (**B**) vs CBD (**C**) males and CBD-exposed females (**F**) vs SHAM (**E**) pups. Each call category was expressed as (number of calls in each category for each subject / total number of calls analyzed in each subject) and represented as the average of each group (SHAM MALE N = 14, CBD MALE N = 7, SHAM FEMALE N = 11, CBD FEMALE N = 11). (**D-G**) CBD modified the vocal repertoire. (**D**) CBD-exposed male emitted more often Short, Trill and Step-up calls, and less Complex Trill and Downward Ramp calls than their SHAM counterparts (**G**). CBD females emitted significantly less Flat, Complex Trill, Downward Ramp, and Inverted-U vocalizations than SHAM (**G**), (SHAM MALE N = 14 light green, CBD MALE N = 7 dark green, SHAM FEMALE N = 11 light orange, CBD FEMALE N = 11 dark orange). Data are represented as Box and whisker plots (first quartile, median, third quartile). Individual data points represent a single animal. Multiple Mann-Whitney *U* test, *p<0.05.

**Table 3:**
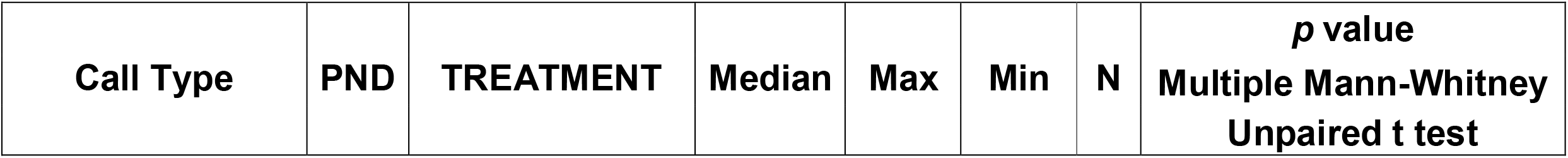
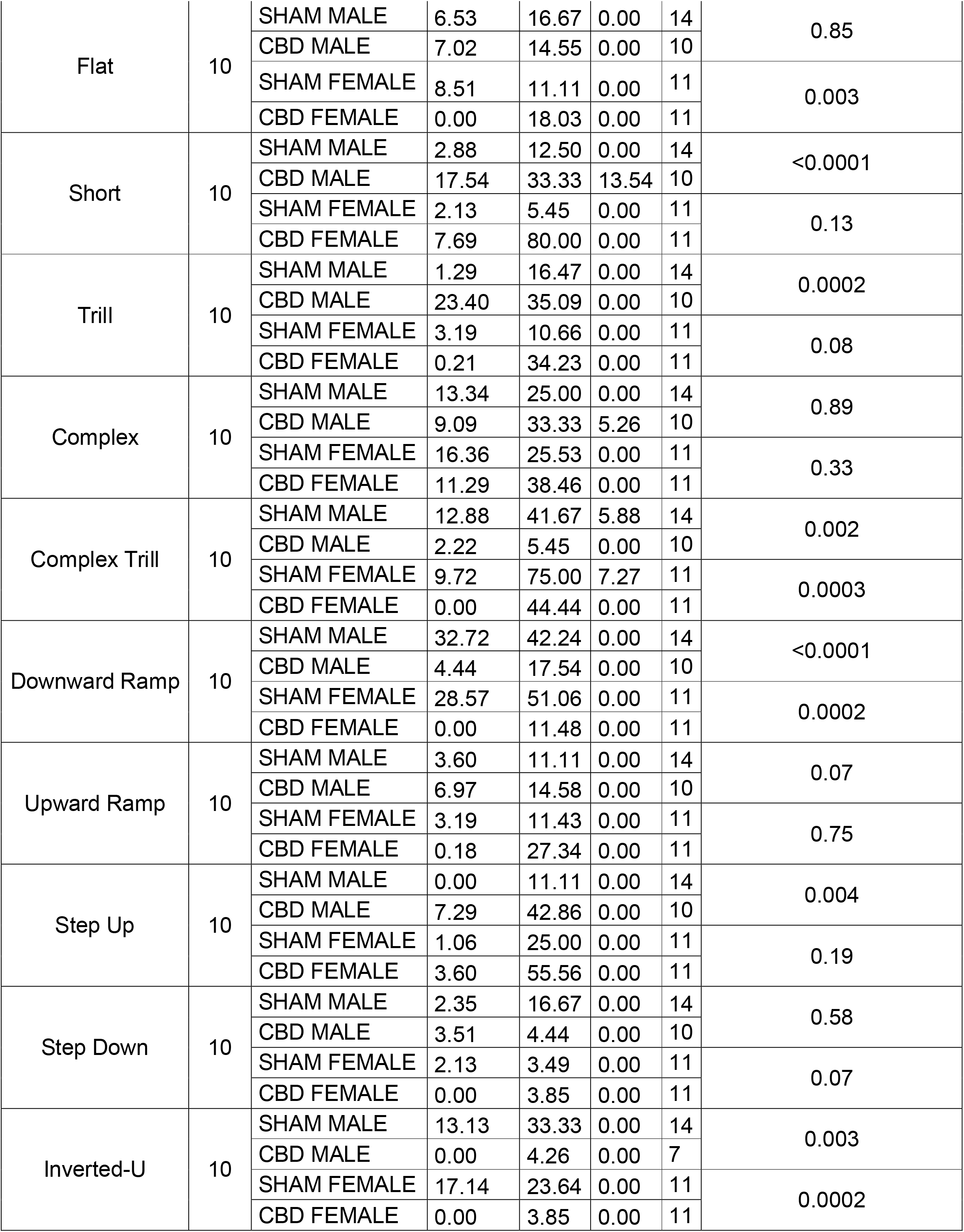
Call type profile is altered in a sex-specific manner in CBD-exposed pups. Data were collected from litters for each condition as described in Methods and Materials. Values are expressed as median, maximum and minimum.

### CBD *in utero* exposure changes the complexity of communication in a sex-specific manner

To test whether the differences found in the syllable repertoire reflect a different development of communication complexity in CBD-exposed pups, we next examined the “call order probability.” Thus, we analyzed the most frequently occurring call combinations (Fig. 4 and Table 4). Certain call sequences were similar in CDB and SHAM males. Indeed, the Inverted-U, Upward ramp, and Complex calls were followed by another Downward Ramp, Upward ramp, and Complex call, respectively, with a similar probability in both CBD and SHAM males (Fig. 4A-B). CBD male showed a significantly lower probability towards the use of Downward Ramp and Complex Trill calls compared to SHAM male pups (Fig. 4C). In the contrast, Trill call was more often used by CBD than SHAM males (Fig. 4C).

**Figure 4.**
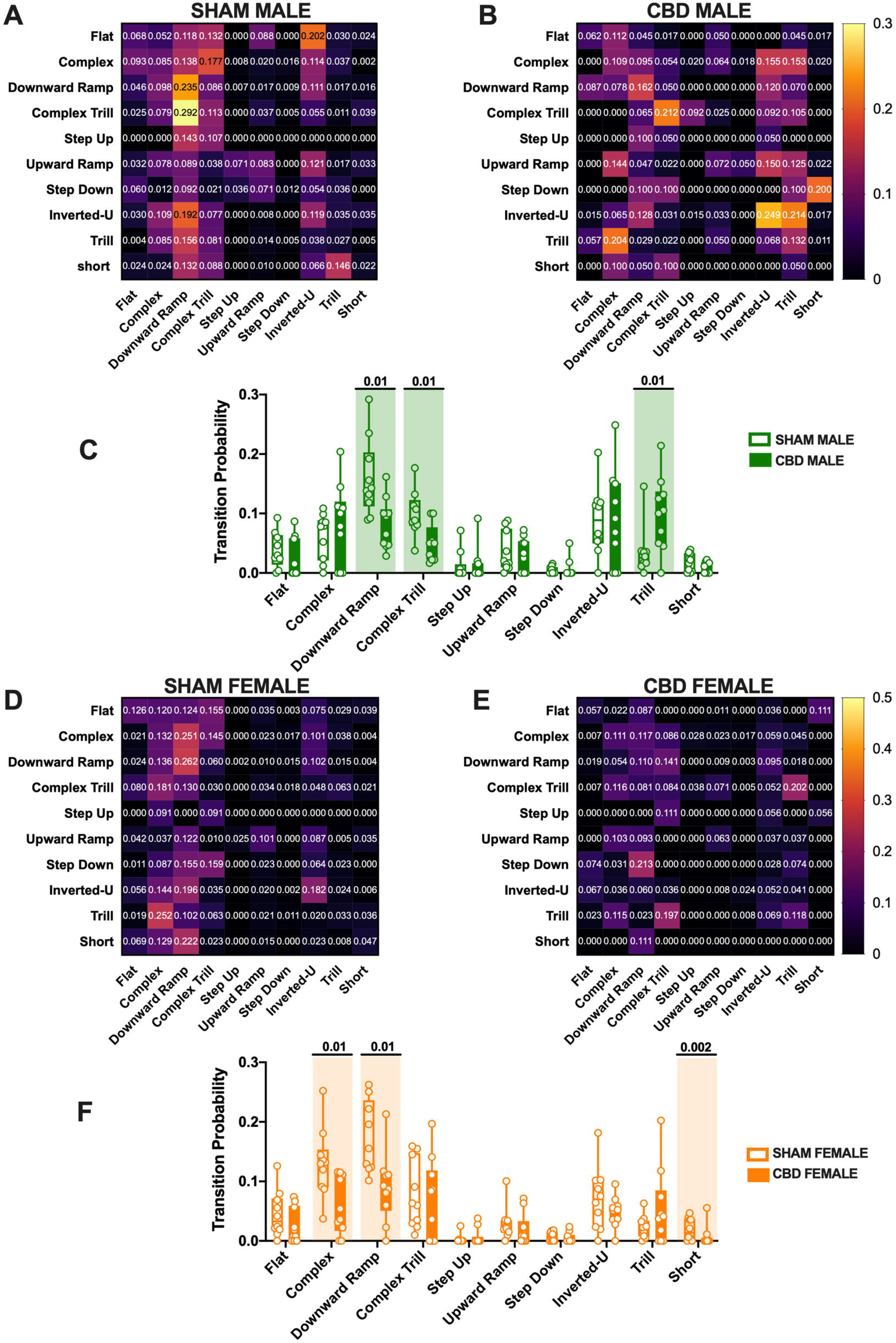
Transitional probabilities for call type transitions within USV bouts in SHAM and CBD pups. Heat maps of transition probabilities in SHAM (**A**) and CBD male (**B**) as well as in SHAM (**D**) and CBD female (**E**) pups. The values in the individual boxes indicate the probability of one call to follow the previous (**A-B-D-E**). The transitional probability was expressed as the mean probability of each transition for each subject. (**C**) The comparison of transitional probabilities shows that CBD-exposed male transitioned significantly less to Downward Ramp and Complex Trill calls than SHAM male pups and that CBD male transitioned more to Trill than SHAM. (**F**) CBD females showed a lower probability to transition to Complex, Downward Ramp and Short calls. Data are represented as Box and whisker plots (first quartile, median, third quartile). Individual data points represent the transitional probabilities of calls emitted by each group. SHAM MALE N = 14, CBD MALE N = 7, SHAM FEMALE N = 11, CBD FEMALE N = 11. Multiple Mann-Whitney *U* test, *p<0.05.

**Table 4:**
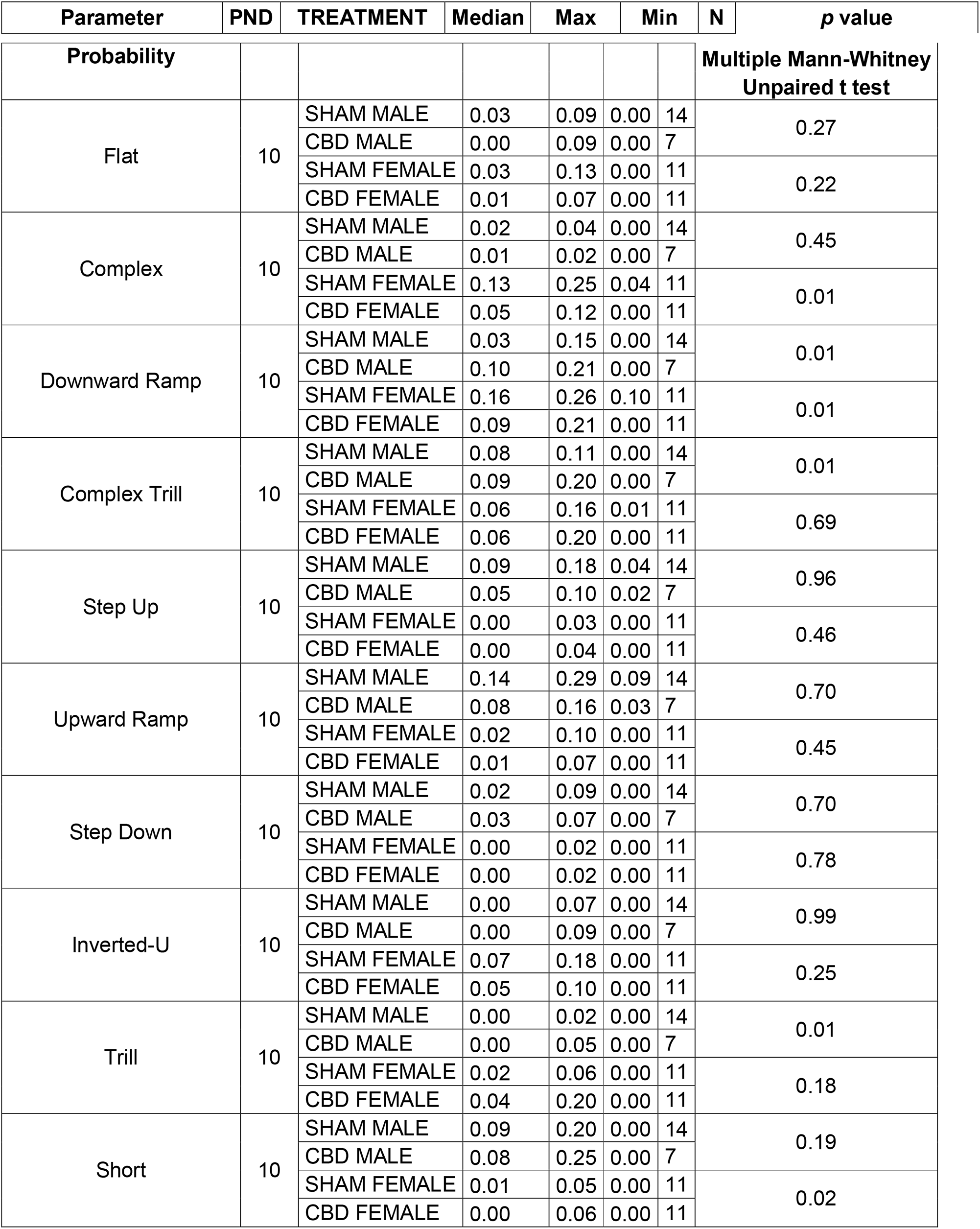
Transitional probabilities for call type transitions within USV bouts in SHAM and CBD pups. Data were collected from litters for each condition as described in Methods and Materials. Values are expressed as median, maximum and minimum.

Finally, we observed a significantly lower probability for the use of Complex, Downward Ramp, and Short calls in CBD females compared to SHAM females (Fig. 4F).

### CBD prenatal exposure impact the motor and discriminative skills during early development in a sex-specific manner

The Homing Test allows the investigation of complex abilities, such as sensory, motor, and odor-detection skills. Thus, we examined homing behavior in PND13, CBD-and SHAM pups (Fig. 5 and Table 5). Gestational CBD significantly reduced the total distance moved (Fig. 5A) only in CBD female pups. Interestingly, these CBD-exposed pups moved slower compared to SHAM female pups (Fig. 5B) and spent less time moving (Fig. 5C) during the 4-min homing test. Moreover, we observed a significant reduction in the distance moved inside the Nest (D) in CBD female pups compared to SHAM pups. On the contrary, no differences were found in the latency to reach, entrances to, and cumulative time spent in the nest, nor in the distance moved in the unfamiliar territory (Fig. 5E-H). Finally, CBD female pups spent more time in the unfamiliar zone, entering more than CBD male pups (Fig 5I-J). These homing test data therefore unveiled an additional sex-specific effect of gestational CBD.

**Figure 5.**
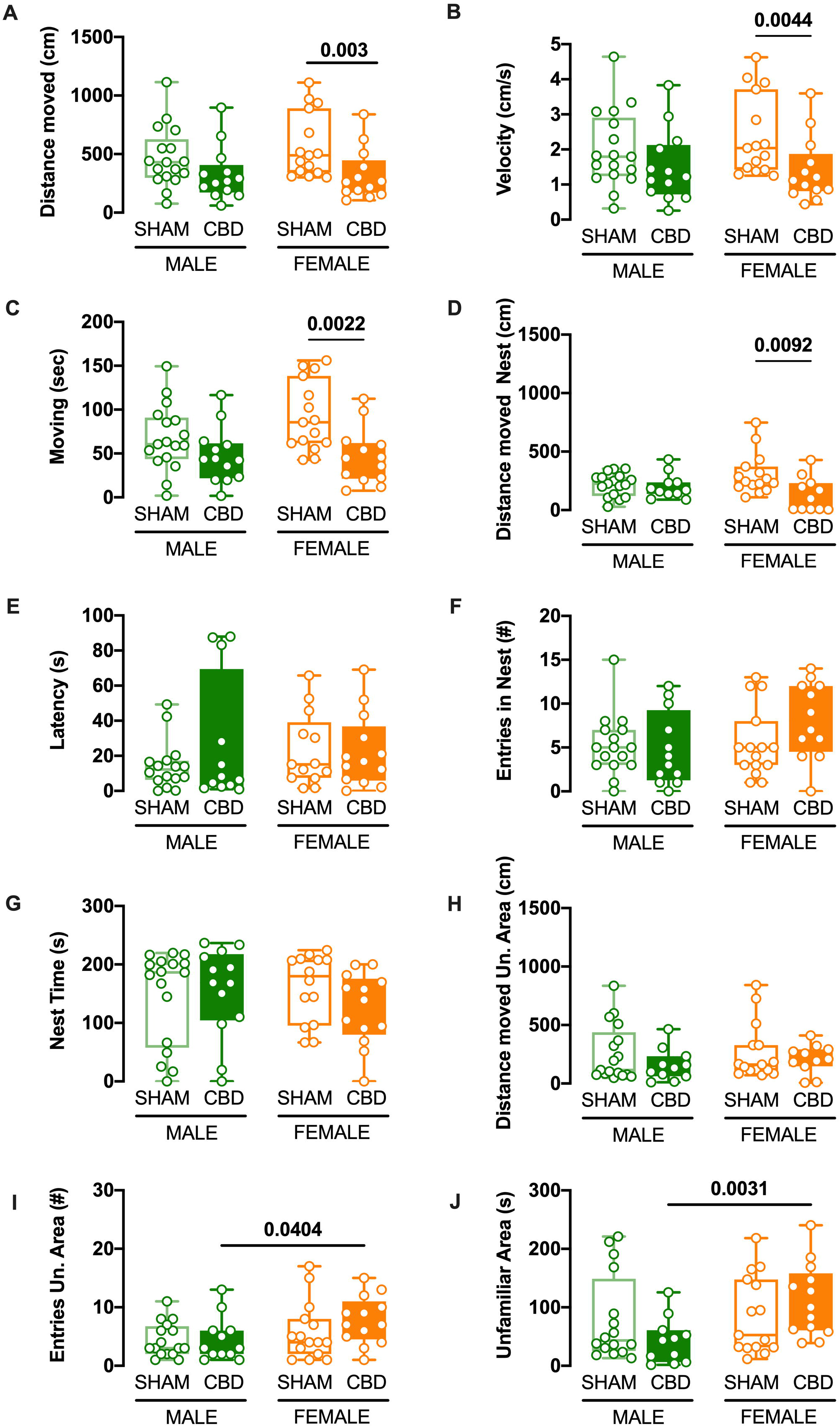
Fetal CBD modifies homing behavior selectively in female pups. (**A-D**) In the female progeny exposed to CBD *in utero*, the total distance moved, the distance moved in the nest, the velocity, and the total time spent moving were diminished. In contrast, these parameters were normal in the male progeny. (**E-H**) Fetal CBD had no discernable effects on the latency to reach the nest, cumulative time spent in the latter, the distance moved in the unfamiliar area or entries into the Nest (**I-J**) CBD females entered more often and spent more time in the unfamiliar zone than the CBD male pups (SHAM MALE N = 15 light green boxplot, CBD MALE N = 13 dark green boxplot, SHAM FEMALE N = 15 light orange boxplot, CBD FEMALE N = 13 dark orange boxplot). Data are represented as Box and whisker plots. Individual data points represent one animal while the line the median. Multiple Mann-Whitney *U* test, *p<0.05.

**Table 5:**
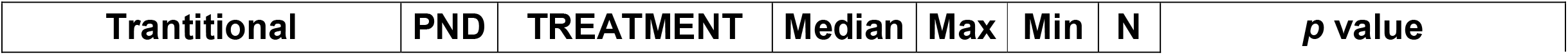
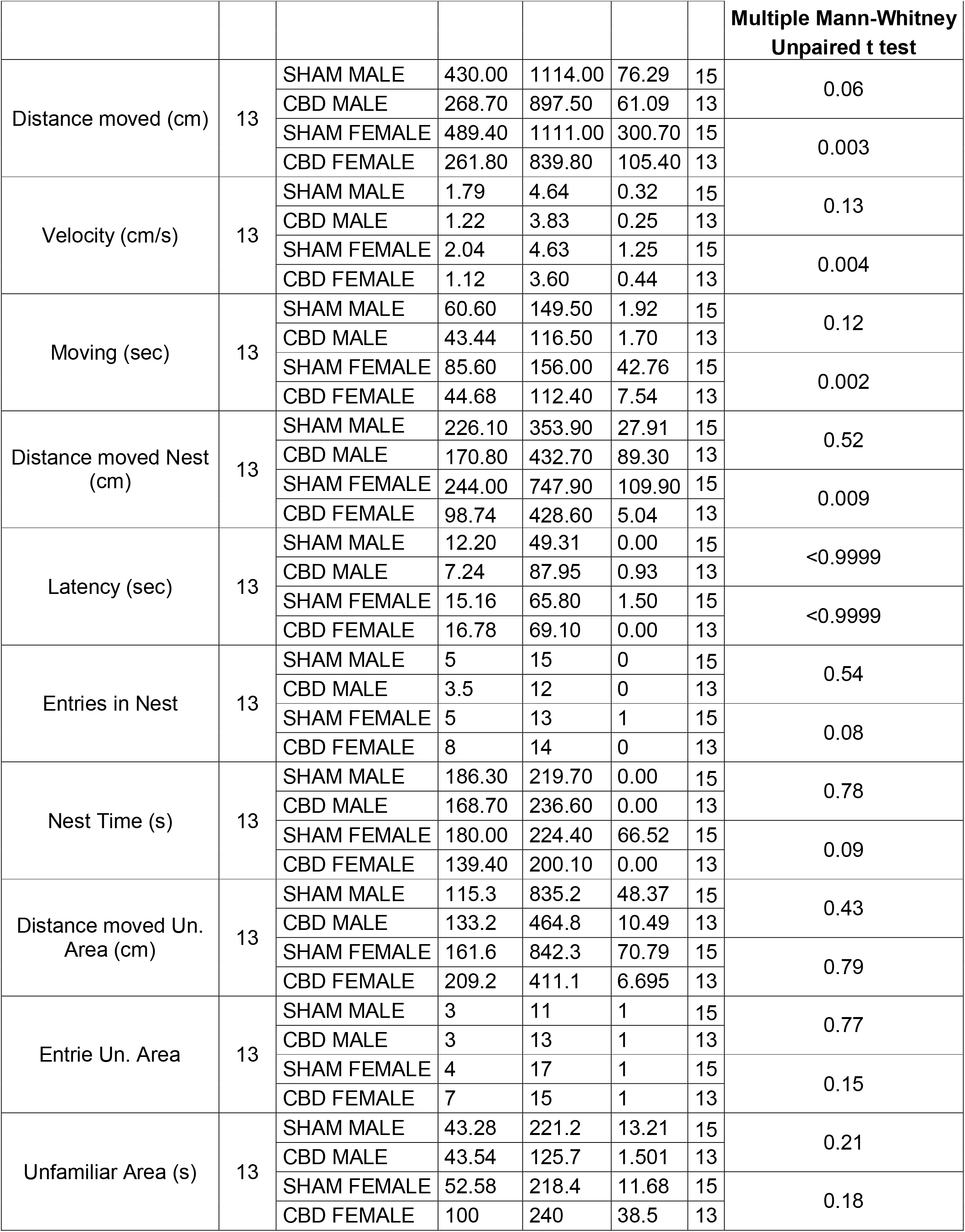
Fetal CBD modifies homing behavior selectively in female pups. Data were collected from litters for each condition as described in Methods and Materials. Values are expressed as median, maximum and minimum.

## Discussion

The consumption of CBD during pregnancy is increasing, but the developmental consequences are still largely unknown. Here, we investigated the sex-specific consequences of prenatal CBD exposure on pre-weaning behaviors. Fetal exposure to a low dose of CBD was associated with increased body weight in male pups during the perinatal period. In addition, the offspring of dams exposed to CBD during gestation showed sex-specific disturbances in their communication, motor skills, and discrimination abilities. Overall, these data indicate that gestational CBD is deleterious to early life behaviors in a sex-specific manner.

In human, gestational cannabis negatively impacts neonatal outcomes (24) and, at birth, the weight of infants exposed to cannabis *in utero* is lower (15,25). Animal models of prenatal THC or synthetic cannabinoid exposure confirm these observations (26). The present results extend these findings to another abundant phytocannabinoid, CBD. We found that maternal exposure to low CBD increases body weight, an effect observed only in males. This sex difference may be linked to differential levels of brain CBD in embryos exposed to CBD (11). In the sole other study that tested *in utero* CBD, no change in the body weight was found in the progeny after weaning or during adulthood (11). This discrepancy may be due to different dam strains (C57BL6/J vs CD1), the time of observation of pups’ growth, or both.

USVs represent one of the earliest markers of neurobehavioral development, allowing quantification of affect, motivation, and social behavior (27–29) USV have an important communicative role in mother-offspring interactions, notably to elicit parents’ attention and care. Thus, understanding mother-pup communication will ameliorate the comprehension and allow the early identification of neurodevelopmental diseases. Pre- and perinatal exposure to psychoactive cannabinoids (e.g., THC) impacts rats’ USVs during infancy in a sex-specific way (23,30). The current results show that *in utero* CBD lowers the number of vocalizing males (not females). CBD-exposed males emitted shorter calls, while CBD-exposed females emitted calls at higher frequency compared to other groups. We found that prenatal CBD also reduced the complexity of the vocal repertoire. Thus, compared to SHAM pups, CBD-exposed pups of both sexes emitted fewer composite calls such as Complex Trill, Downward Ramp, and Inverted-U, when compared to their SHAM littermates. In addition, male CBD-exposed pups employed monosyllabic calls (e.g., “Short” calls) more often than other groups. Interestingly, we found that the same categories of calls found altered in CBD-treated animals reflected lower probabilities to use those calls, suggesting a call-specific effect of gestational CBD. The complexity of the vocal repertoire increases during life (31) and though the precise meaning of these vocalizations remains unclear, one could hypothesize that prenatal exposure to CBD changes early communication skills. Our observation is reminiscent of altered ultrasonic communication reported in several murine models of autism (i.e., fmr1^y/−^, BTBR, Shank1^−/−^) (32–35) and is in line with human studies showing abnormal cry characteristics in sick toddlers with diseases affecting the central nervous system, including autism spectrum disorders (36). Thus, it is tempting to conclude that communication deficit is a common and early marker of neurodevelopmental diseases. Homing behavior requires sensory and cognitive skills (e.g., to differentiate the scent of the original cage) as well as motor skills (e.g., to navigate to the original litter). CBD had sex-specific effects on homing; only CBD-treated females showed an overall reduction in motor activities (i.e., distance traveled, speed, and total time spent moving). In addition, CBD-treated females entered the unfamiliar area more often and spent more time in the unfamiliar area than CBD-treated male pups, suggesting differential development of sensory and cognitive abilities.

Taken together, this study reveals sex-specific cognitive impairments in early life associated with fetal CBD. This work challenges the view that CBD is a universally safe compound and warrants further study of the developmental consequences of prenatal CBD.

## Author contributions

Daniela Iezzi: Conceptualization, Data curation, Formal analysis, Validation, Writing— original draft, review and editing.

Alba Caceres: Data curation, Formal analysis.

Pascale Chavis: Conceptualization, Supervision, Methodology, Writing—, review and editing.

Olivier JJ Manzoni: Conceptualization, Supervision, Funding acquisition, Methodology, Project administration, Writing— original draft, review, and editing.

The authors declare no conflict of interest.

## Acknowledgements

The authors are grateful to the Chavis-Manzoni team members for helpful discussions and to Dr. A.F. Scheyer for critical reading and help with writing the manuscript.

